# A temporal hierarchy of object processing in human visual cortex

**DOI:** 10.1101/2023.03.07.531635

**Authors:** Rui Xu, Wenjing Zhou, Xin Xu, Zhipei Ling, Hesheng Liu, Bo Hong

## Abstract

Our brain constructs increasingly sophisticated representations along the ventral visual pathway to support object recognition. To understand how these representations unfold over time, we recorded human intracranial electroencephalography responses to 120 object images at five consecutive stages of the ventral pathway from V1 to occipitotemporal areas. Using representational similarity analysis, we confirmed that response patterns were more strongly driven by low-order stimulus properties at early stages and high-order category information at late stages, respectively. Interestingly, response patterns also became less stimulus-driven and more categorical at all stages over time, from ∼100 to 200 ms post-stimulus. During this period, we found significant noise correlation between single-trial response patterns across stages, indicating tight inter-areal coupling. Thus, multi-areal recurrent processes may be essential in building high-order object representations.

## MAIN TEXT

Primates can extract abstract information about objects (e.g., category and identity) from their visual forms in a split second^1^. This important function of object recognition is supported by the ventral visual pathway^2^ extending from the primary visual cortex (V1) in the occipital pole rostrally to the inferotemporal cortex (IT)^3^ in monkeys and occipitotemporal cortex (OTC)^4,5^ in humans. Neural responses represent increasingly higher-order information ascending the pathway, from simple spatiotemporal filters in V1 to category and identity in IT/OTC, and responses at the late stages along the pathway show tolerance^6–8^ to image transformations that alter low-order but not high-order features.

Although the spatial hierarchy is well established, much less has been known about the temporal profile of responses in the ventral visual areas. First, previous studies found IT/OTC responses represent different information at various latencies, e.g., category vs. identity^9–11^ and individual parts vs. their configuration^12^, which indicates involvement of intra- and inter-areal recurrent processing^13^. However, these studies typically used a limited set of stimuli, e.g., faces and artificial shapes, and it is unclear if their findings generalize to other objects. Second, early visual areas such as V1 were implicated in both feedforward and recurrent modes of visual processing^14^. Still, the temporal dynamics in these areas during object recognition and how they relate to the dynamics in IT remain poorly understood.

## Results

### Spatiotemporally resolved object responses

To understand how object representations unfold across areas and over time, we examined high gamma activity^15,16^ of human intracranial electroencephalography (iEEG) responses recorded from electrodes that collectively covered much of the cerebral cortex (n = 1227; fig. S1A). Subjects performed a one-back task viewing serially presented images of clear, isolated objects for 300 ms every 900 to 1050 ms while maintaining fixation. The stimuli were 120 *instances* of objects from three *categories*: faces, places, and other objects (Fig. 1A), which all correspond to representative category-selective regions in human OTC^17^. The rationale is that the dissimilarity relations between neural response patterns to various objects at a given area and time should depend on which information is encoded there and then^18,19^. Therefore, we didn’t normalize low-order features across categories; in addition, we divided each category into two subcategories or *classes* and ensured that same-class instances shared certain low-order feature such as contrast or silhouette (Fig. 1A). In this design, both low-order features and category information differ at the class level, and more importantly they predict distinct structures of neural dissimilarity across instances, which will later help distinguish low- and high-order neural representations.

**Fig. 1.**
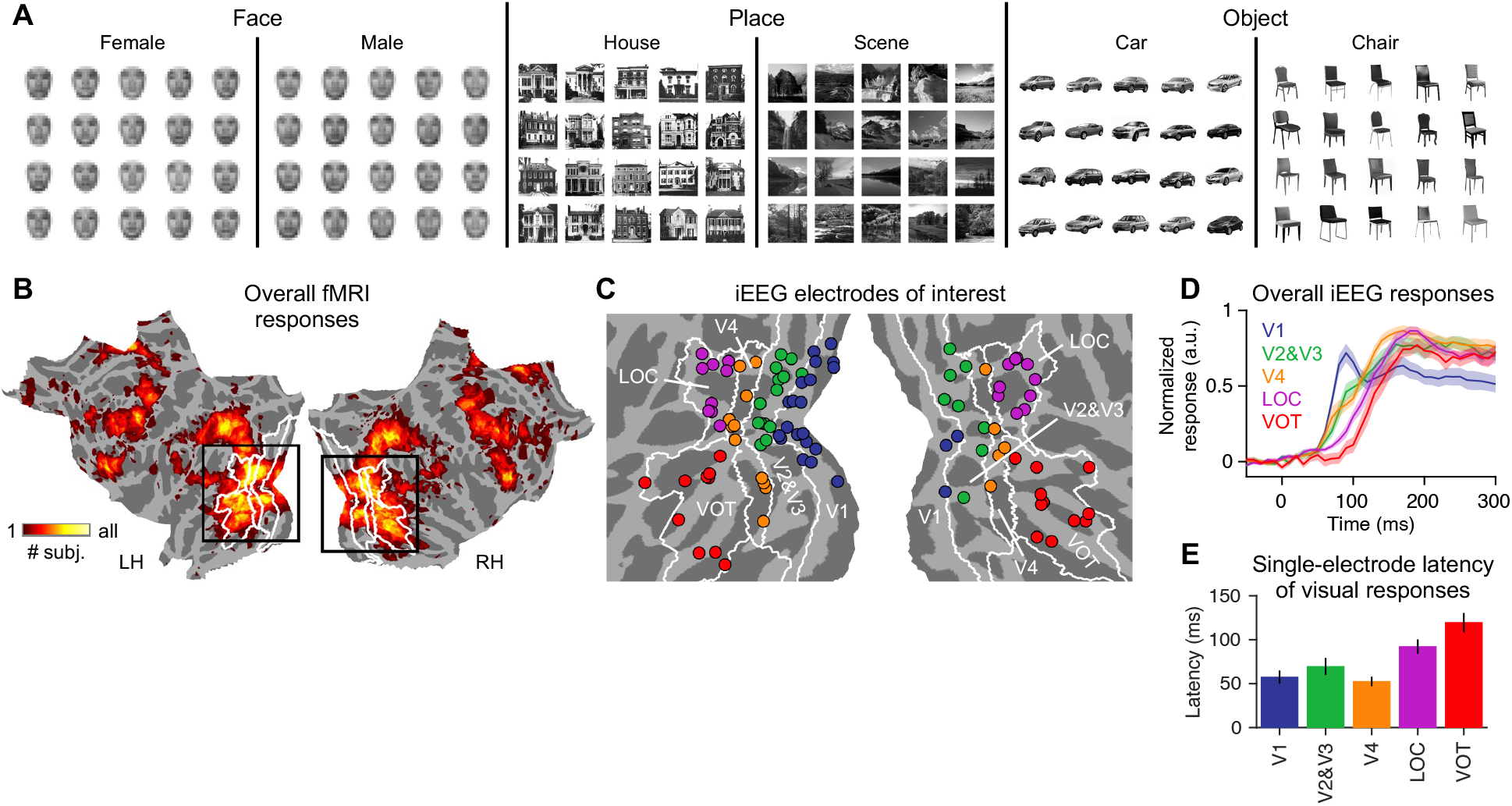
Stimuli and iEEG recordings. (**A**) The stimuli are 120 images of object instances belonging to 6 classes and 3 categories. We pixelated the face images for privacy. (**B**) The number of subjects in which a vertex showed significantly positive fMRI responses to the stimuli (P < 10^−4^), shown as an overlay on flattened surfaces of fsaverage. White contours are boundaries of ventral visual areas. Black rectangles match the field of view in (C). (**C**) Electrodes of interests on fsaverage colored by area. White contours are the same as in (B). (**D**) The overall iEEG responses to all stimuli. Colors indicate the area, as in (C). (**E**) Cumulative distribution of single-electrode latency of visual responses.

We searched for iEEG electrodes showing positive responses (50-300 ms post-stimulus) to visual stimuli and above-chance class-level decoding (n = 122; see Methods). The majority of them (n = 101) were in ventral visual areas and co-localized with positive functional MRI (fMRI) responses to the same stimuli (Fig. 1B and figs. S1A-S1B), which were the focus of this study. They covered five consecutive stages of the ventral pathway (Fig. 1C and fig. S1C) according to a population atlas^20^, including V1 (n = 23), V2&V3 (n = 22), V4 (n = 16), and two parts of OTC^4,5^, i.e., lateral occipital cortex (LOC; n = 19) and ventral occipitotemporal cortex (VOT; n = 21). The overall iEEG responses followed separable time courses across stages (Fig. 1D), and the latency of responses was significantly shorter in V1 through V4 than LOC and VOT (paired T-test, P < 0.01 for V1 vs. LOC and V2&V3 vs. VOT, P < 0.001 for V4 vs. LOC, P < 10^−4^ for V1 vs. VOT and V4 vs. VOT; Fig. 1E), which is consistent with feedforward signal propagation and validates the spatiotemporal resolvability of iEEG.

### Time course of object representations across areas

The criteria for selecting the electrodes of interest ensured that their responses differed by class. A closer look showed that the relative amplitude of responses to different classes often changed substantially over time (see Fig. 2A for examples), which implies that the same area may represent different information over time. However, it is inherently limited to characterize the neural representation of objects, which is multidimensional, based on univariate response profiles of individual channels. Thus, we used representational similarity analysis instead. As the first step, we built 120 × 120 representational dissimilarity matrices (RDMs; Fig. 2B and fig. S2E) for each area and time point separately by calculating a modified version of standardized Euclidean distance of multi-electrode response patterns across instances (see Methods). To better visualize the structure of RDMs, we used multidimensional scaling (MDS) to approximate inter-instance neural dissimilarity with Euclidean distance (Fig. 2C). Non-random RDMs emerged earlier in V1 than VOT, and the relative layout of instances by class differed between areas and transformed over time within each area.

**Fig. 2.**
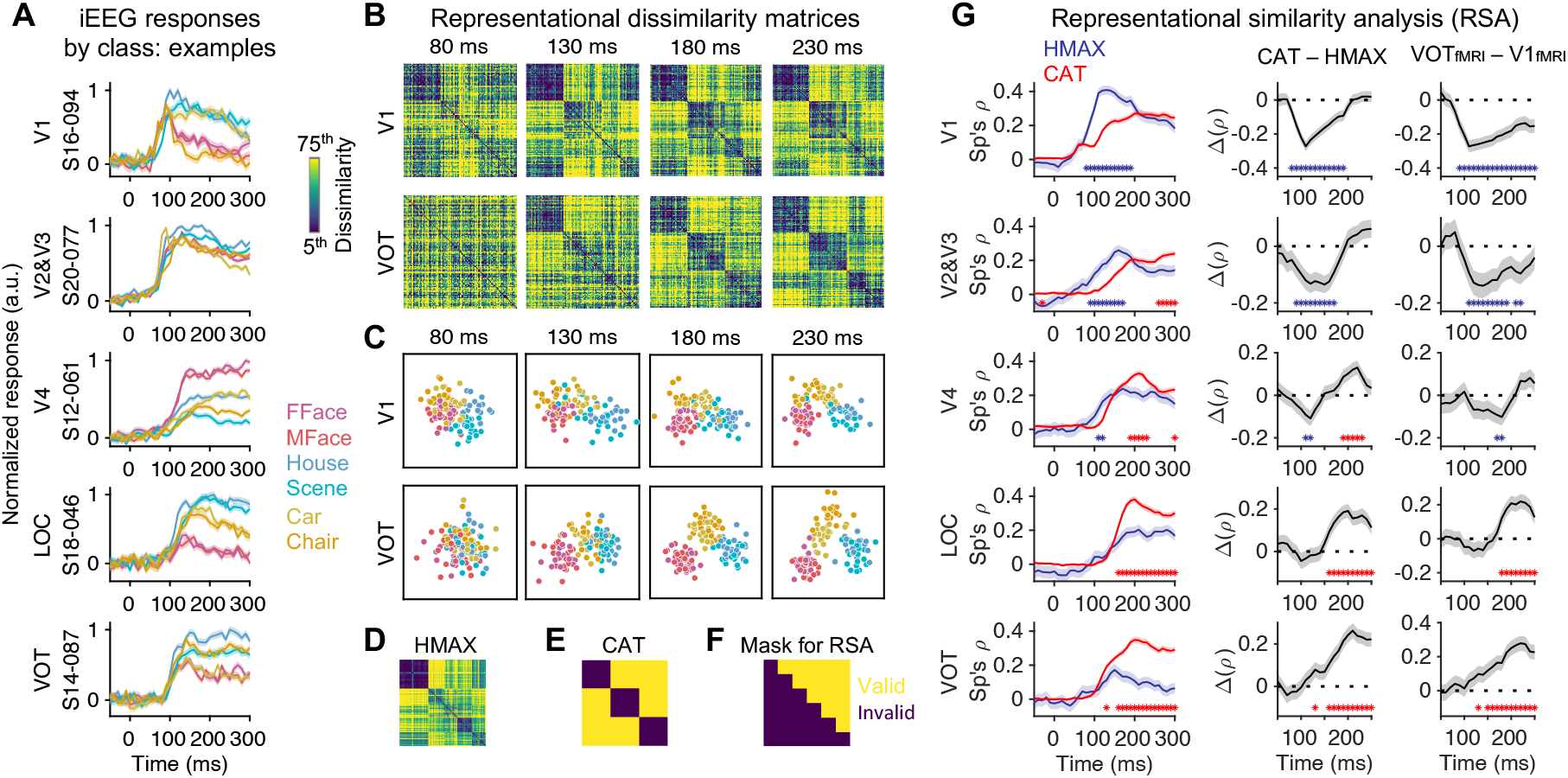
Spatiotemporal dynamics of object representations. (**A**) Neural responses by class at −50 to 300 ms relative to stimulus onset in example electrodes. Solid lines and shadings denote the mean and standard error of mean over same-class trials. (**B**) RDMs derived from iEEG responses in V1 and VOT at several time points.Colormap covered the 5^th^ to 75^th^ percentile of dissimilarity values in each RDM. (**C**) MDS visualization of iEEG RDMs in (B). Instances are shown as dots and colored by class. (**D**) As (B), reference RDM derived from the output of HMAX-C2 units (HMAX). (**E**) As (D), reference RDM of ideal categorical representation (CAT). (**F**) RDM entries used to calculate representational similarity. Valid (used) and invalid (discarded) entries are indicated by color. (**G**) Left, representational similarity (Spearman’s ρ) of iEEG vs. reference RDMs. Mid, the difference between the representational similarity of iEEG vs. CAT (high-order reference) and iEEG vs. HMAX (low-order reference). Right, as mid, with RDMs derived from fMRI responses in VOT and V1 as high and low references, respectively. Solid lines and shadings denote mean and standard error of bootstrap distribution. Blue/red asterisks indicate when similarity vs. low/high-order reference is significantly higher (p<0.05).

We next compared the neural RDMs to two reference RDMs derived from HMAX^21^, a classic model of low-level vision, and ideal categorical (CAT) representation (Figs. 2D and 2E; see Methods). A neural RDM corresponding to low- and high-order processing should be more similar to HMAX and CAT references, respectively. Because same-class instances by design both shared low-order features and belonged to the same category, we calculated representational similarity (Spearman’s ρ) using only different-class dissimilarities (Fig. 2F) to better distinguish low- and high-order neural RDMs. The results were consistent with the spatial hierarchy of object processing. The neural RDMs of V1 at many time points were significantly more similar (p<0.05) to the low-order (HMAX) reference than the high-order (CAT) one. Most LOC and VOT RDMs showed an opposite pattern. V2&V3 and V4 RDMs showed an intermediate pattern, of which some were closer to the low-order reference and others the high-order one (Fig. 2G, left). Interestingly, in all five areas, the neural RDMs over time became less similar to the low-order reference and more similar to the high-order one, and the difference of similarity to the two references (CAT – HMAX) increased during ∼100 to 200 ms post-stimulus (Fig. 2G, mid). Thus, object processing becomes more sophisticated not only from the early to late stages of the ventral pathway but also from early to late time points at each stage, i.e., there is a time-domain parallel of the spatial hierarchy for object vision.

The observation of spatial and temporal hierarchies didn’t depend on the choice of reference RDMs. First, the pattern of results didn’t change (Fig. 2G, right and fig. S3) when we instead used RDMs derived from fMRI responses in V1 and VOT (fig. S2D; see Methods) as low- and high-order references, respectively, as in previous studies^22,23^. Second, we also found correspondence^24–27^ between bottom/top layers of deep convolutional neural networks (DCNNs; fig. S2F; see Methods) and neural RDMs at early/late stages along the ventral pathway and early/late time points (fig. S3). The RDMs of top DCNN layers didn’t match high-order neural RDMs unless we used the SHINE operation^28^ to equalize low-order features before feeding the stimuli into the DCNNs, and the best-matching DCNN layer after SHINE still didn’t match the neural RDMs as well as fMRI, which are consistent with the findings that DCNNs don’t fully account for the behavioral and neural responses^27,29,30^. We noted that an alternative method^24,31^ based on predicting neural responses with DCNN outputs was unsuitable for assessing brain-DCNN correspondence in this study (fig. S4; see Methods), as it failed a crucial test of validity^24^ using our data. Finally, we derived the same spatial and temporal hierarchies without specifying any references by relating the neural RDMs across areas and time. RDMs of different areas were, in general, more similar/dissimilar if they were close/distant in time (fig. S5), and the transformations of RDMs from early to late time points and from early to late areas were largely along the same direction (fig. S6).

### Time course of tolerance to image transformations across areas

A hallmark of high-order object processing is tolerance to image transformations. To quantify this, we trained class-level decoders using iEEG responses to normal stimuli and tested them using responses to normal or size-reduced stimuli. We were able to collect data for this analysis from one subject, as we needed electrode coverage of multiple areas and additional trials. We divided the electrodes into two groups, V1 to V3 and V4 to VOT (n = 11/3; Fig. 3). Decoding accuracy of 1/3-sized stimuli reached significance (p < 0.05) slightly later than normal (150 vs. 120 ms) in V4 to VOT, consistent with the latency reported in a previous human iEEG study^6^. The 1/3-sized accuracy also reached significance in V1 to V3, which was unexpected, but much later than normal (180 vs. 40 ms). Besides a longer latency, the performance drop of 1/3-sized vs. normal was bigger in V1 to V3, indicating weaker tolerance. Thus, responses at the early stages of the ventral pathway were initially not tolerant to image transformations but became tolerant around 200 ms, consistent with the temporal hierarchy we derived independently using representational similarity analysis. The decoding accuracy of 2/3-sized stimuli was close to normal (latency = 40/130 ms in V1-to-V3/V4-to-VOT), suggesting the magnitude of size change fell short of breaking stimulus-driven class separability.

**Fig. 3.**
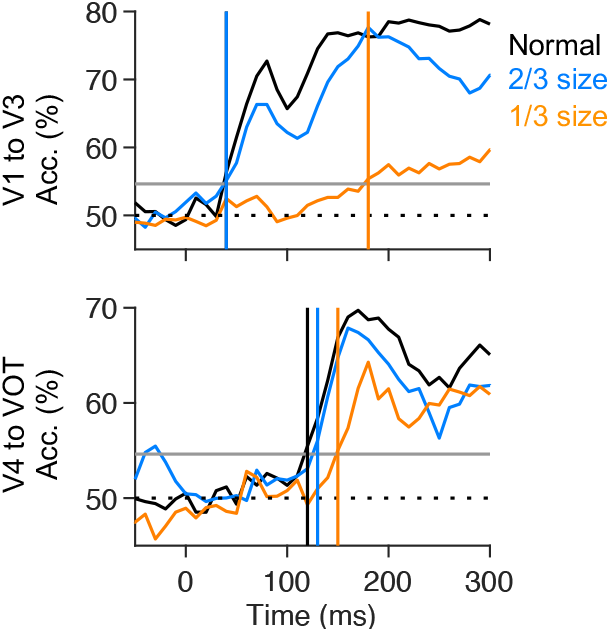
Size-tolerance decoding accuracy. Top, V1 to V3; bottom, V4 to VOT. Color indicates the image size of test trials in decoding analysis. Dashed and solid gray lines indicate the chance level and permuted threshold (P < 0.01), respectively. Vertical lines indicate decoding latency for different sizes.

### Trial-by-trial association of response patterns between areas

The increase in processing order began around the onset of responses at the latest stages of the ventral pathway (Fig. 1D), and it occurred for an extended period and concurrently at all stages. Therefore, the temporal hierarchy we found may result from non-feedforward processes, including recurrence across stages^32–34^ and a common top-down input^35^. In either case, we expect shared trial-by-trial neural variability between areas^36^. To quantify this (see Methods), we converted single-trial multi-electrode patterns to univariate responses along object-decoding axes, and then computed the noise correlation (Spearman’s ρ) between converted responses across areas of the same subjects. In six subjects with electrodes in both V1 to V3 and V4 to VOT, the correlation of responses across areas but at the same time (Fig. 4A) was significantly (p < 0.05) higher than baseline at multiple time points between 100 and 200 ms, which confirmed that there was shared trial-by-trial variability between areas. We further examined cross-area, cross-time correlation (Fig. 4B). The lack of consistency in the electrode coverage across subjects may contribute to the variations in their correlation patterns. Nonetheless, we found a clear trend in the subject-averaged pattern that the correlation was stronger between adjacent than distant time points (r = −0.87, p = 1.8 × 10^−139^), suggesting a dynamic inter-areal coupling. The data doesn’t favor either V1 to V3 or V4 to VOT as the “driver” of their shared variability, as there wasn’t a clear lead/lag between them.

**Fig. 4.**
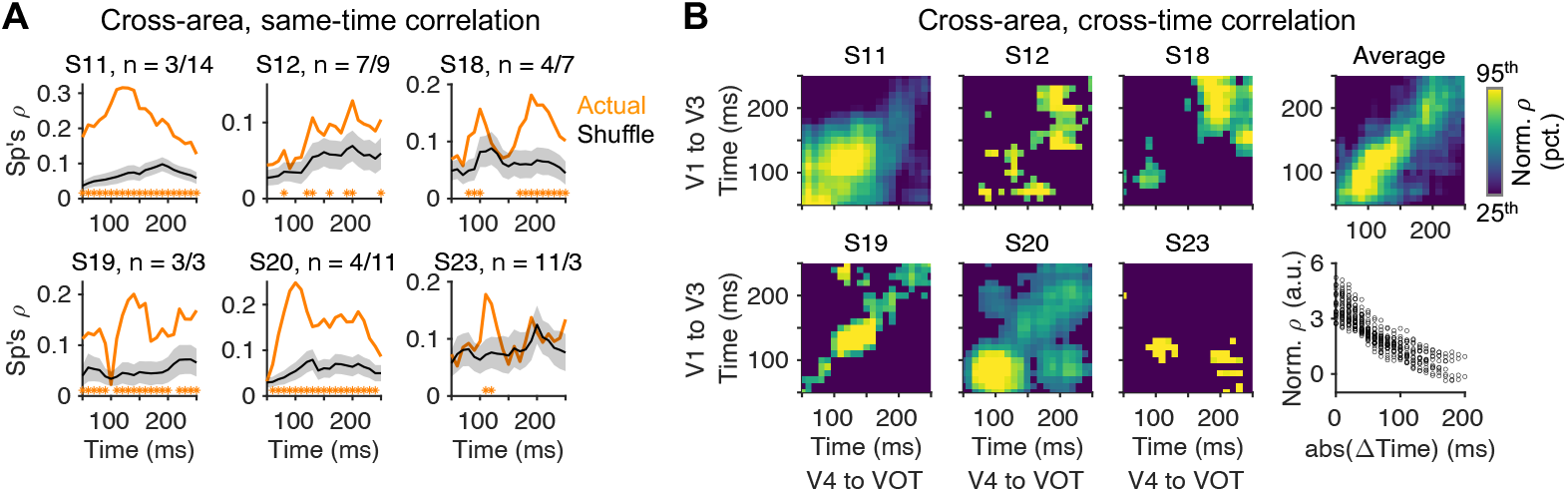
Trial-by-trial association across areas. (**A**) Cross-area correlation (Spearman’s ρ) of concurrent multielectrode patterns projected to common axes. Orange/black, correlation derived from actual/shuffled data. Shadings indicate standard error of null distribution (shuffle). Orange asterisks indicate when the correlation is significantly higher than shuffle (P < 0.05). n indicates # electrodes in V1 to V3 and V4 to VOT per subject, respectively. (**B**) Cross-area, cross-time correlation matrices of individual subjects (left) and population average (right-top). X-axis and Y-axis indicate the time of V4 to VOT and V1 to V3 when calculating correlation, respectively. Normalized ρ is shown (actual divided by shuffle). We set insignificant correlation values to 0 before averaging .Right-bottom, population-average correlation vs. inter-areal delay (absolute value). Circles indicate entries of the correlation matrix.

## Discussion

Using human iEEG, we provided a spatiotemporally resolved account of object representations in the ventral visual pathway: going from early to late time points between ∼100 and 200 ms in every area of the pathway, we found higher-order representational dissimilarity structures and better tolerance to image transformations, as one would expect going from early to late areas along the pathway. Previous studies^22,23^ reported correspondence between magnetoencephalography (MEG) representations of objects at early/late time points and fMRI representations in early/late visual areas. The temporal hierarchy we found is consistent with such correspondence but cannot be deduced from it since MEG and fMRI only resolve either space or time but not simultaneously^37^. For instance, a similar pattern of MEG-fMRI correspondence would have been found if a low-order representation was established in V1 before a high-order one in OTC, and neither evolves once established.

Despite the prevalence of non-feedforward connections in the primate visual system^38^, a prominent view^3^ holds that object recognition is supported by largely feedforward mechanisms, and an important piece of evidence for this view is that IT/OTC responses as early as ∼100 ms support object decoding^6,7^. In contrast to this view, we found substantial representational changes in OTC after its initial responses, including being driven less by low-order features and more by object categories related to OTC’s spatial organization^17^, and they were concurrent with similar changes in lower areas as well as tight coupling between OTC and lower areas. The results suggest an important role of recurrence in extracting object category information by the human brain, even though purely feedforward mechanisms like DCNNs can readily perform this task. Recurrence has been strongly implicated when recognition is challenging due to blur, clutter, occlusion, etc., as extra processing time is needed^23,30,39–41^. Our findings are broadly consistent with such observations, while support yet less conditional involvement of recurrence using clear, isolated stimuli. Future causal experiments^35,42–45^ guided by connectional anatomy^38,46^ may help discern between different types of recurrence, including the interaction between OTC and lower areas, top-down feedback^35,47–49^, and intra-areal recurrence.

The mechanism of recurrent object processing may not be inherently more complex than other visual functions known to rely on recurrence, such as contour integration^32^. In principle, downstream neurons at a later stage of the visual hierarchy may build a coarse presentation at first based on their initial inputs, which may then help upstream neurons at an earlier hierarchical stage with smaller receptive fields disambiguate^50^ what they see, and the disambiguated signals may, in turn, contribute to a refined downstream presentation, and so on. A similar mechanism with different sites of downstream and upstream neurons may underlie object recognition, and its further characterization likely requires fine-grained stimulus design combined with novel computational models^34^.

## ACKNOWLEDGEMENTS

We thank Chen Song and Dan Zhang for their assistance in data collection. This work was supported by the National Key R&D Program of China (2017YFA0205904 to B.H.) and the National Natural Science Foundation of China (NSFC 62061136001 to B.H.).

## AUTHOR CONTRIBUTIONS

Conceptualization: R.X., B.H. Methodology: R.X., B.H. Investigation: R.X., B.H., W.Z., X.X., Z.L. Visualization: R.X. Funding acquisition: B.H. Project administration: B.H., R.X., W.Z. Supervision: B.H. Writing – original draft: R.X. Writing – review and editing: R.X., H.L., B.H.

## DATA AND MATERIALS AVAILABILITY

The data of this study will be de-identified and deposited in a public repository before publication.

## METHODS

### Participants and paradigm

Twenty-one patients (7 female, 12-47 years old) with pharmacologically intractable epilepsy participated in this study, intracranially implanted with iEEG electrodes to localize seizure foci. They were admitted to Tsinghua University Yuquan Hospital (TYH; n = 13) or Chinese PLA General Hospital (PGH). The Institutional Review Boards at both hospitals and Tsinghua University approved the study. We obtained informed consent in writing from all subjects before participation. All subjects had normal or corrected-to-normal vision.

Subjects viewed 120 grayscale square images of clear, isolated objects from three categories: faces, places, and other objects. Each category was further divided into two subcategories or classes (six in total), i.e., female faces, male faces, houses, scenes, cars, and chairs, and there were 20 instances per class. We ensured that same-class instances shared some visual features such as orientation and silhouette. We used the stimuli in a previous fMRI study^51^. In each trial of the main iEEG experiment, an image pseudorandomly drawn from the full set was shown for 300 ms in the center of a black screen at ∼6.5 visual degrees and then disappeared for 600-750 ms. At the same time, a small white fixation point was continuously present in the center of the screen. For four subjects (2 electrodes of interest) recruited at the beginning of the study, images were shown for 500 ms, followed by blanks of 750-1000 ms. In a control experiment to assess tolerance to image size, images were presented at full, 2/3, or 1/3 of the regular size for 200 ms, followed by blanks of 700-850 ms. We instructed subjects to maintain their fixation and press a key if they saw the same image (regardless of size) shown in the previous trial, i.e., to perform a 1-back task (P_1-back_ = 12%). We put 45 to 60 trials in a block, and subjects could rest between blocks. We collected 348 to 1566 trials per subject for the main experiment and 870 trials in one subject (S23; 14 electrodes of interest) for the control experiment.

Thirteen subjects also participated in a modified version of the main experiment in an fMRI session consisting of 3 to 6 runs. We used a blocked design for optimal contrast-to-noise ratio. Each run had six 32-sec stimulus blocks, one for each class, interleaved with seven 16-second fixation blocks. Each stimulus block consisted of 20 or 40 trials showing same-class images in a pseudorandom order, for 300 or 600 ms, followed by 500-ms or 1000-ms banks. We used the version with fewer, longer trials at the beginning of the study (n = 6), and the fMRI response patterns didn’t differ systematically between versions. Images were presented in the center of a white screen at ∼10 visual degrees with small jittering of position. Subjects performed the same 1-back task. We conducted both iEEG and fMRI experiments using Psychtoolbox-3^52^.

### Data acquisition and preprocessing: iEEG

Subdural surface grids/strips and/or depth electrodes were implanted based on clinical considerations only. The iEEG signals were sampled at 1200 Hz and 512 to 1024 Hz using g.Tec and Nicolet amplifiers at TYH and PGH, respectively, along with a high-pass filter (cutoff = 0.1 Hz) and a notch filter at 50 Hz (to remove power line noise). Four electrodes placed on the external surface of the skull with the contacts facing away from the skull were used as ground and reference (two electrodes each). We manually identified iEEG electrodes showing epileptic discharge and removed them from further analyses. We then removed electrodes and trials showing other artifacts semi-automatically using *ft_rejectvisual* in FieldTrip. We also discarded the first two trials of a block and trials in which the image repeated the previous trial or subjects pressed the key. We ended up with 211 to 1150 valid trials per subject in the main experiment and 548 in the control experiment (used in Fig. 3 only).

In this study, we focused on high gamma or broadband activity, which is a non-oscillatory component with no particular frequency peak and correlates with neuronal spiking ^53,54^. We set 10-ms spaced time points from −200 to 800 ms (relative to the stimulus onset) as time points of interest (TOIs), and for each TOI, we built ten four-cycle Morlet wavelets and had their center frequency logarithmically spaced from 110 to 140 Hz. The spectral bandwidth of the wavelets together covered 82.5-185 Hz, and the wavelet duration was between 9.1 and 11.6 ms. We computed power spectral density (PSD) for each wavelet. We took the logarithm to make it more Gaussian, which was then normalized (subtraction before division) for each electrode separately, based on the mean and standard error of log(PSD) between −100 and 0 ms over trials and experiments. Finally, we averaged log-normalized PSD over wavelets and used it as the metric of iEEG responses in subsequent analyses. For visualization in Fig. 1D, we averaged the metric over all trials and then divided it by the max value across all time points for each electrode separately. For Fig. 2A, we averaged the metric over same-class trials and then divided it by the max value across all time points and classes.

### Data acquisition and preprocessing: MRI

We collected structural (MPRAGE; TR/TE = 2530/3.45 ms; 1 mm isotropic) and, in some cases, blood-oxygenation level-dependent (BOLD) functional (EPI; TR/TE = 2000/30 ms; voxel size = 3.125 × 3.125 × 4 mm^3^; 25 slices; FA = 90 degrees; no gap) MRI images in a Philips Achieva 3.0 T TX scanner before subjects were implanted. Using FreeSurfer^55^, individual brain surfaces were reconstructed based on structural MRI images, nonlinearly transformed to a template brain (fsaverage), and parcellated according to a population atlas^20^. FreeSurfer also generated volume-based parcellation (e.g., of cortical areas, subcortical structures, and white matter) of individual brains. Using FreeSurfer functional analysis stream (FS-FAST), volume-based fMRI data processing steps included motion correction, slice timing correction, and quadratic linear detrending. The volume-based data were then projected onto individual surfaces and smoothed (FWHM = 5 mm). A general linear model computed each class’s regression coefficient (beta). We also computed an omnibus contrast (stimulus vs. fixation). We transformed the statistics to fsaverage for further analyses.

### Electrode localization and selection

We followed a semi-automatic pipeline^56^ with reported precision within 3 mm to localize electrodes in the brain. We manually labeled the location of electrodes based on postoperative computer tomography (CT) images. We aligned the CT images to same-subject structural MRI images via rigid transformation using a mutual information-based algorithm and manual adjustment if needed. To address brain deformation due to the insertion of electrodes, a constrained energy-minimization algorithm was used to pull surface electrodes onto smoothed pial surface (pial-outer-smoothed) while minimizing the displacement of electrodes and deformation of grids/strips. For depth electrodes, we considered them cortical if they were closer to cortical than subcortical voxels and the distance to the nearest cortical voxel was no larger than 3.5 mm. Finally, for all surface and depth cortical electrodes, we found the nearest vertex on the individual pial surface and the corresponding vertex on fsaverage surfaces.

We narrowed the cortical electrodes down to the electrodes of interest for further analyses in three successive steps (fig. S1), using data from the main experiment. First, we looked for visually responsive electrodes by comparing mean responses over TOIs of 50 to 300 ms vs. −100 to 0 ms (fixation) for trials of each class separately. We kept electrodes showing significantly stronger-than-fixation responses (paired T-test, P < 10^−4^) to at least one class. All P values reported in this study are two-sided. Second, we performed single-electrode, class-level decoding analysis and kept electrodes showing significantly above-chance-level accuracy (permutation test, P < 0.05). Lastly, we discarded electrodes not in ventral visual areas according to the population atlas or in a vertex where at least one subject showed significantly positive fMRI responses (omnibus contrast, uncorrected P < 10^−4^).

We set single-electrode latency of visual responses as the earliest TOI (between 10 and 300 ms), starting from which five consecutive TOIs showed significantly (paired T-test, P < 10^−4^) stronger-than-fixation (−100 to 0 ms) responses to at least one class.

### Single- and multi-electrode decoding analysis

For single-electrode decoding analysis, we used linear SVM classifiers (LIBSVM; C = 1) to decode the object class of single trials, taking responses from 50 to 300 ms (26 TOIs) as features. We built 15 sets of binary classifiers, one for each pair of classes. For all class pairs and electrodes, we chose the same number of trials (30 per class, 60 in total) pseudorandomly from all valid trials in each cross-validation (CV) fold, of which we used 2/3 and 1/3 to train/test the classifier. 30 was the minimal number of total valid trials for any subject and class. We normalized training and test data in each CV fold by subtracting the mean and dividing by the standard error of training data for each TOI separately. We did 50 CV folds per classifier for each electrode and reported decoding accuracy as the mean over 15 class pairs (chance level = 50%). We performed a permutation test to assess statistical significance by shuffling the class label of all valid trials and then running decoding analysis as normal (15 classifiers, 50 CV folds per classifier). We did 2000 permutations per electrode to construct a robust null distribution. We considered decoding accuracy significant (P < 0.05) if it was larger than the 97.5^th^ percentile of the null distribution.

We performed size-tolerant multi-electrode decoding analysis for each TOI and area separately, taking multi-electrode response pattern at a given TOI and area as features. We determined the significance and latency of decoding similarly to single-electrode decoding, with a stricter threshold (P < 0.01). In each CV fold, we drew 45/23 trials per class and size (2/3 of the minimal number of total valid trials for any class and size) from main/control experiments as training/test data. We shuffled the joint class/size label of trials in the main and control experiments separately for the permutation test. The earliest TOI (between 10 and 300 ms), starting from which five consecutive TOIs showed significant decoding accuracy, was taken as decoding latency.

### Representational similarity analysis

Because each instance was presented to a varying number of electrodes, we used a modified version of standardized Euclidean distance to quantify the inter-instance dissimilarity of multi-electrode response patterns. First, we converted responses to up to 120 instances (averaged over same-instance trials) to the Z-score for each electrode separately. Second, we computed the dissimilarity between the two instances as the root mean square of the difference between Z-score vectors consisting of electrodes to which both instances were presented. We built one 120 × 120 representational dissimilarity matrix (RDM) of iEEG for every area and TOI (Fig. 2B and fig. S2E). To visualize the RDMs, we applied non-classic MDS using *mdscale* in MATLAB with ten dimensions to get low-stress solutions. We showed the first two MDS dimensions and used Procrustes transformation to stretch/stress and rotate the 120-instance clouds to align them across areas and time (Fig. 2C).

The 120 × 120 RDM of HMAX (Fig. 2D and fig. S2A) was computed as standardized Euclidean distance across rows of 120 × # units matrix consisting of the output of all units at the C2 layer of HMAX model, using *pdist* in MATLAB (*Distance = ‘seuclidean’*). The 120 × 120 RDMs of DCNNs (fig. S2F) were computed in similar ways based on the output of units in six selected layers (each) of two representative DCNNs, which were, from bottom to top (L1 to L6), ‘pool1’, ‘pool2’, ‘pool5’, ‘fc6’, ‘fc7’, and ‘fc8’ in Alexnet, and ‘max_pooling2d_1’, ‘activation_10_relu’, ‘activation_22_relu’, ‘activation_40_relu’, ‘avg_pool’, and ‘fc1000’ in Resnet-50, respectively, following a recent study^27^. We created two sets of RDMs per DCNN, one using the images presented to subjects as input (‘Raw’), the other using the same images but with some low-order features equalized with the SHINE (spectrum, histogram, and intensity normalization and equalization) operation^28^ as input (‘SHINE’). The 120 × 120 RDM of ideal categorical (CAT) representation (Fig. 2E and fig. S2B) was built by setting all same-category entries as zero (similar) and the rest as one (dissimilar).

For fMRI, we started with 6 × 6 inter-class RDMs and then converted them to 120 × 120 inter-instance RDMs. For each area, we pooled vertices from both hemispheres of a subject that showed significantly positive responses (omnibus contrast, P < 10^−4^) to build a 6 × # vertices matrix of beta value. We then computed standardized Euclidean distance across rows of the beta matrix. The results were averaged over subjects with at least 200 significant vertices in the area to generate the inter-class RDM, in which all diagonal entries were zero. Before averaging, we normalized individual subjects’ RDMs with a division by the square root of the number of significant vertices. Finally, we built the corresponding inter-distance RDM (fig. S2D) by setting dissimilarity between two instances as that between respective classes of the instances.

When computing representational similarity (RS) between inter-instance RDMs, we only considered 6000 above-diagonal (for the RDMs were all symmetrical) entries corresponding to the dissimilarity between instances from different classes (Fig. 2F and fig. S2C). We didn’t consider dissimilarity between same-class instances because it was undefined for fMRI RDMs and expected to be small for both low- and high-order representations due to our choice of stimuli. To assess statistical significance, we created a bootstrap distribution of RS using iEEG RDMs derived from a subset of data pseudorandomly drawn from all trials with replacement. We made 2000 rounds of bootstrap, and trials were selected once per round for all electrodes of the same subject. We considered an iEEG RDM significantly more similar to one reference RDM than another (P < 0.05) if the RS of the former reference was higher in more than 97.5% of bootstrap rounds.

Because iEEG RDMs were noisy, especially those of early TOIs, when comparing iEEG RDMs with each other, we used 6 × 6 inter-class RDMs built the same way as inter-instance RDMs but based on responses averaged over same-class trials. To avoid bias toward a stronger correlation between same-area, close-time RDMs, we calculated RS between inter-class RDMs based on two separate halves of trials (split-half RS). In fig. S5, we did bootstrap resampling and calculated split-half RS in each bootstrap round (500 rounds, one split-half per round) to assess statistical significance. In fig. S6, where a significance test wasn’t needed, we averaged RS over 50 split-halves of the full dataset for less noisy results.

### Predicting single-electrode responses with DCNN output

We tested how well single-electrode, instance-level response can be modeled as linearly mixed output of units in selected layers of DCNNs taking raw or SHINED stimuli as input (see “Representational similarity analysis”). We used partial least squares (PLS) regression^24^ (# components = 5) to estimate the mixing weights based on responses to some instances and used the learned weights to predict responses to other instances. Changing # components didn’t alter the pattern of results (data not shown). The performance of prediction was measured as the cross-validated normalized mean square error (norm. MSE) using *goodnessOfFit* in MATLAB (*cost_func = ‘NMSE’*), which is effectively 1 – goodness-of-fit (r^2^). A value of 0 indicates a perfect fit, 1 indicates fit as good as a flat line through the mean response, and larger than 1 indicates a bad fit likely due to overfitting in training. We did two types of six-fold cross-validation, and in each fold, we either used 5/6 and 1/6 of instances, regardless of their class, as training and test data, respectively (object generalization) or used instances of 5 class as training data and the other class as test data (class generalization).

### Reliable class- and instance-level modulation of single-electrode responses

We quantified class- and instance-level modulation of single-electrode, single-trial responses based on the partition of sums of squares (SS): the total sum of squares (SS_Total_), i.e., the sum of squares of deviation from single-trial response to the total mean summed over all trials, equals the sum of SS_Class_ (deviation from class mean to total mean), SS_Instance_ (deviation from instance mean to class mean), and SS_Error_ (deviation from single-trial response to instance mean). We only considered instances presented in at least three trials for each electrode to separate SS_Instance_ from SS_Error_. SS_Class_ and SS_Instance_ divided by SS_Error_ (normalized SS) can be used as class- and instance-level modulation metrics, respectively.We didn’t separate class-level modulation within and across categories. Because non-error SS terms are always overestimated with limited data due to noise, we created a control term for SS_Class_ (SSC_Class_) by shuffling the class label of instances without changing the instance label of trials. We created another control term for SS_Instance_ (SSC_Instance_) by shuffling the instance label of trials within each class. We subtracted the SSC terms (mean of 500 shuffles) from respective non-error SS terms and added both to SS_Error_ when calculating normalized SS.

### Trial-by-trial association of multi-electrode response patterns

To measure the within-subject, trial-by-trial association between multi-electrode response patterns, we projected them to axes perpendicular to classification hyperplanes of the 15 binary classifiers, which are comparable across areas and time. In each CV fold of decoding analysis, we used 30/15 trials per class as training/test data in five subjects with coverage of both V1 to V3 and V4 to VOT and 20/9 trials in another subject with fewer trials available. Unlike other decoding analyses, here we applied the trainer classifier to test trials of all classes. We computed Spearman’s ρ using projected responses along each of the 15 axes for each class to avoid spurious correction since classes modulated responses. To address the additional confounding effect of instance-level modulation, we constructed a baseline by shuffling the order of same-instance trials before computing ρ. We didn’t analyze trials of instances presented fewer than three times to a subject. We assessed statistical significance based on the null distribution constructed from 2000 shuffles.

**Fig. S1.**
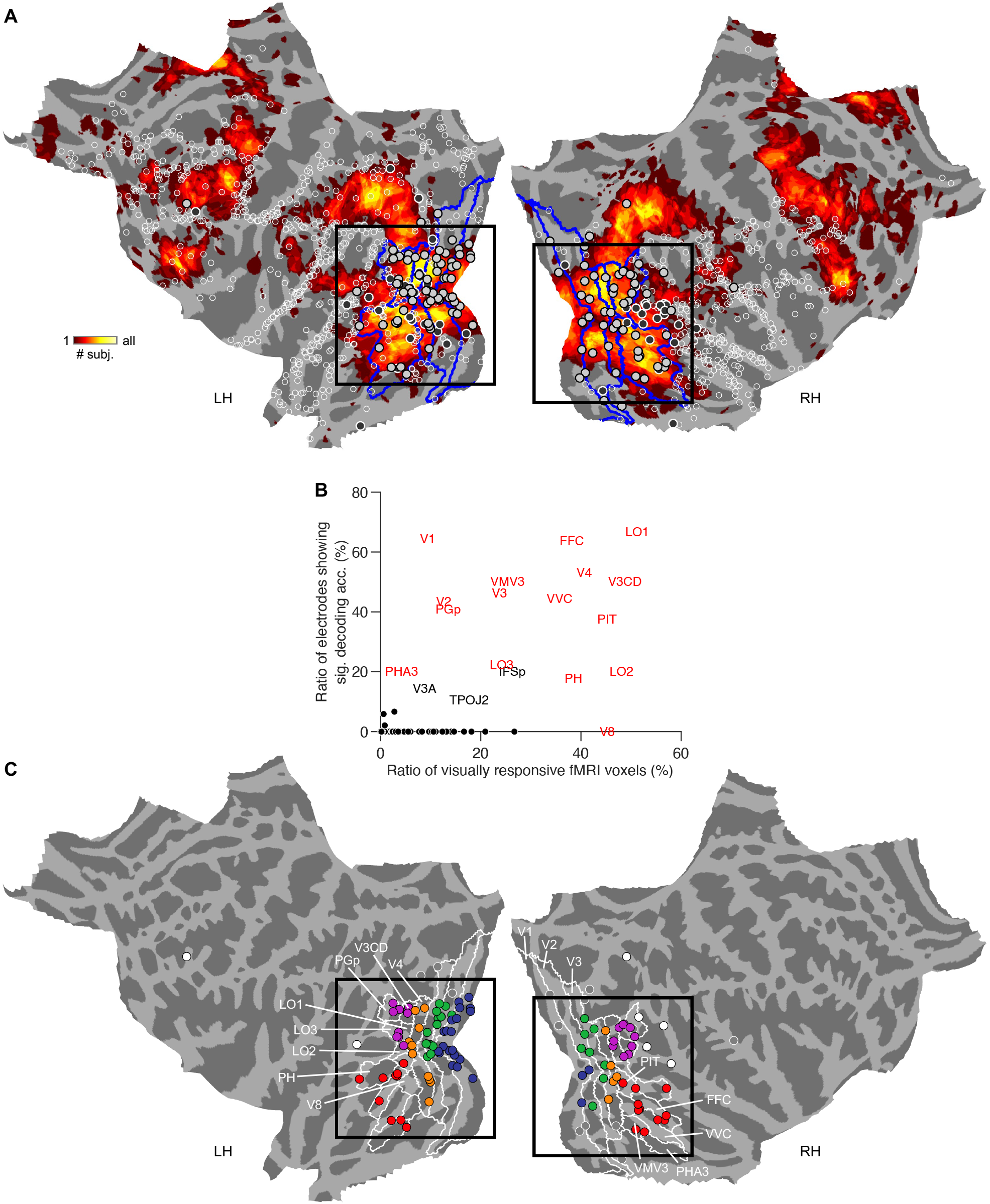
Coverage and grouping of iEEG electrodes. (**A**) All cortical electrodes on flattened surfaces of fsaverage, along with fMRI responses to the same stimuli shown as in Fig. 1B. Hollow circles, electrodes not showing significantly stronger-than-fixation responses (P < 10^−4^) to any object class (n = 1055, including four cortical electrodes not shown due to cuts made to flatten the surface). Gray/black circles, visually responsive electrodes showing / not showing significantly above-chance-level decoding accuracy (P < 0.05; n = 122/50). Overlay indicates # subjects showing significantly positive fMRI responses (P < 10^−4^). Blue contours are boundaries of five areas/stages along the ventral visual pathway (V1, V2&V3, V4, LOC, and VOT), many of which consist of multiple areas in a population atlas^20^. Black rectangles match the field of view in Fig. 1C. Most decodable electrodes were found in ventral visual areas and co-localized with strong fMRI responses. (**B**) The ratio of decodable electrodes vs. visually responsive fMRI voxels in the same area (both hemispheres combined) for all cortical areas in the atlas covered by at least five electrodes. We show ventral visual areas as red text, areas outside the pathway where>10% of electrodes were decodable as black text, and the other areas as black dots. iEEG decodability and fMRI visual responsiveness were significantly correlated (r = 0.65, p = 2.8 × 10^−10^), suggesting good consistency between modalities. (**C**) Decodable electrodes on flattened surfaces of fsaverage. Colored and white circles indicate electrodes in one of the five ventral visual areas (electrodes of interest) and the other cortical areas, respectively (n = 101/7), both located in a vertex in which at least one subject showed significantly positive fMRI responses (P < 10^−4^). Gray circles indicate electrodes in a vertex not showing significantly positive fMRI responses in any subject (n = 14). White contours are boundaries of ventral visual areas in the atlas (see labels in white text).

**Fig. S2.**
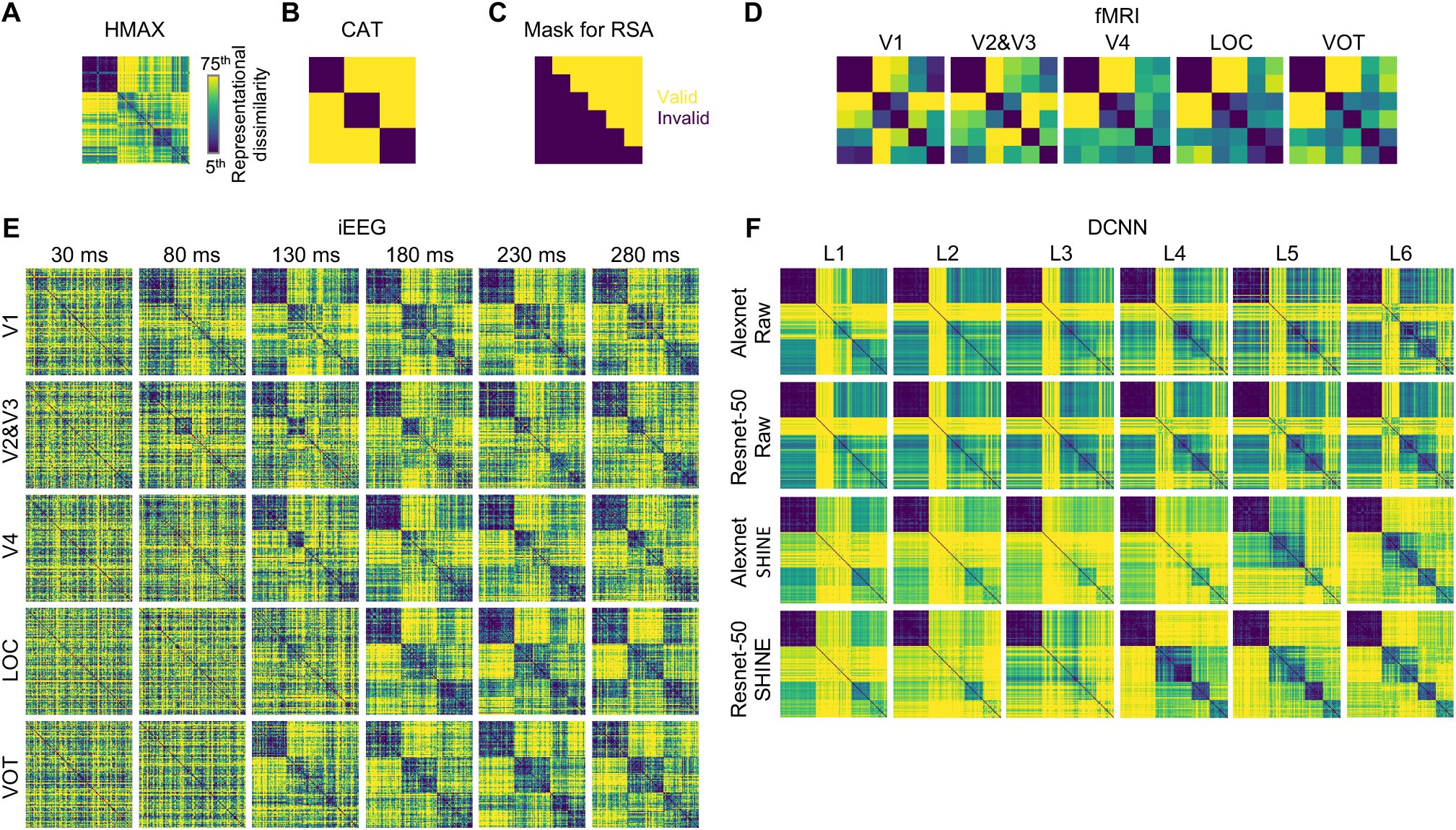
Representational dissimilarity matrices of all modalities. (**A**) RDM derived from the output of HMAX-C2 units (HMAX). Colormap covered the 5^th^ to 75^th^ percentile of dissimilarity values in the RDM. (**B**) As (A), of ideal categorical representation (CAT). (**C**) RDM entries used to calculate representational similarity. Valid (used) and invalid (discarded) entries are indicated by color. (**D**) As (B), derived from fMRI responses in different areas. (**E**) As (B), derived from iEEG responses in different areas at several time points. (**F**) As (B), derived from out of layers L1 to L6 in Alexnet and Resnet-50 using raw or SHINED stimuli as input. RDMs of L1 through L6 were very similar using raw stimuli, but not so much using SHINED stimuli: RS of L1 vs. L6 of Alexnet/Resnet-50 was 0.79/0.58 and 0.26/−0.02, using raw and SHINED stimuli as input, respectively. L6-SHINE was also distant from L1-raw (RS of Alexnet/Resnet-50 = 0.12/0.03) but much closer to fMRI-VOT (RS = 0.51/0.59) than was L6-SHINE (RS = 0.00/−0.15).

**Fig. S3.**
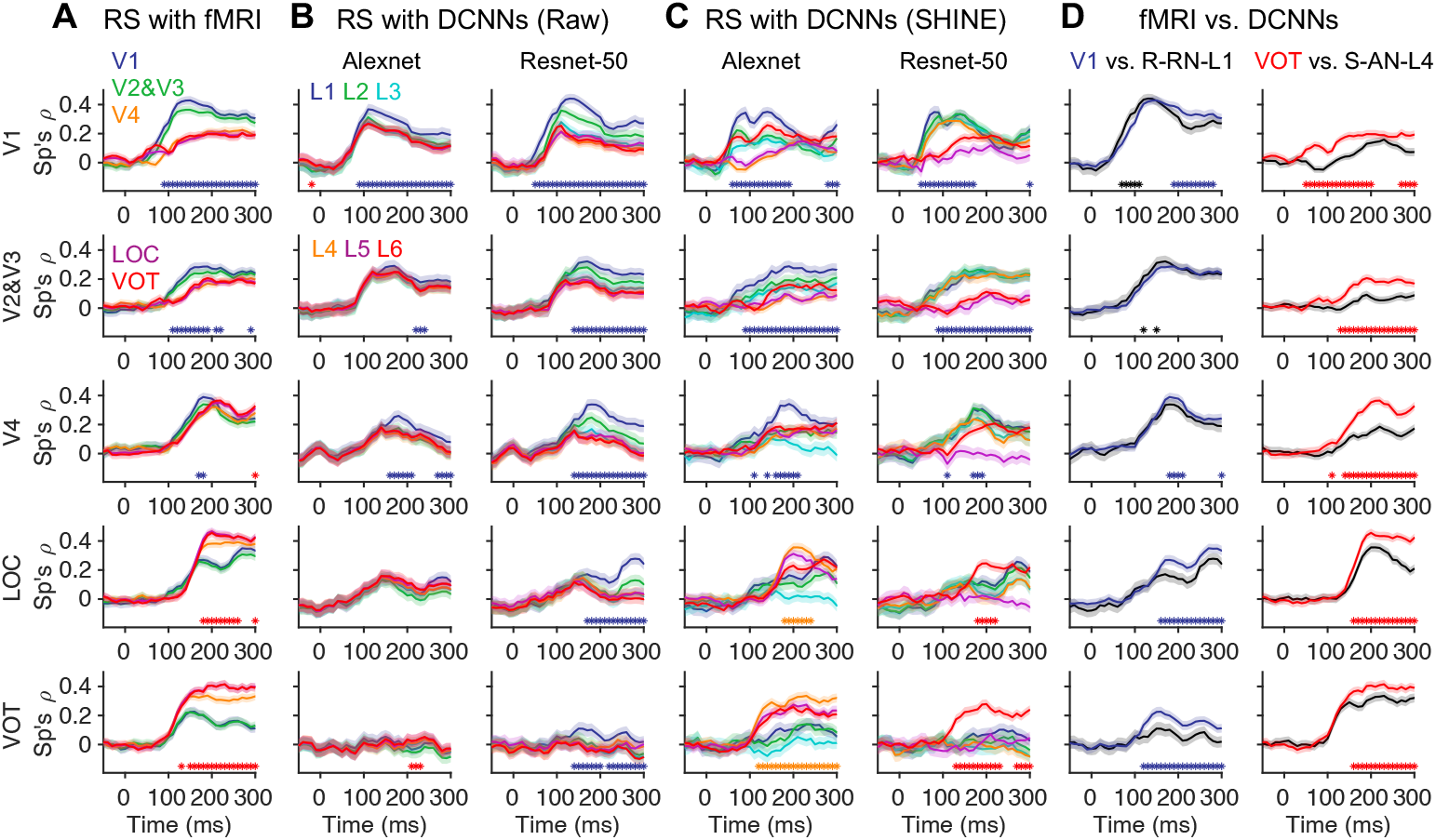
Relating representational dynamics to fMRI and DCNNs. (**A**) Representational similarity (RS) of iEEG vs. fMRI. Solid lines and shadings denote mean and standard error of bootstrap distribution. Blue/red asterisks indicate when RS_fMRI-V1_/RS_fMRI-VOT_ is significantly higher (p<0.05). For iEEG RDMs across all areas, their RS vs. fMRI-V1 was similar to RS vs. fMRI-V2&V3, and RS vs. fMRI-V4, RS vs. fMRI-LOC, and RS vs. fMRI-VOT were similar to each other. The iEEG RDMs were, in general, more similar to same-area RDMs: iEEG-V1/V2&V3 were close to fMRI-V1/V2&V3, iEEG-LOC/VOT were close to fMRI-V4/LOC/VOT, and iEEG-V4 were similarly close to all fMRI RDMs. For iEEG RDMs of the same area, their RS vs. fMRI-V1/V2&V3 peaked earlier than fMRI-LOC/VOT. (**B**) As in (A), RS of iEEG vs. layers L1 to L6 of Alexnet (left) and Resnet-50 (right) taking raw stimuli as input. Blue/red asterisks indicate when RS_L1_/RS_L6_ is significantly higher. (**C**) As in (B), with DCNNs taking SHINED stimuli as input. For Alexnet, L4 was the best-matching high-order reference for iEEG among the layers, and blue/orange asterisks indicate when RS_L1_/RS_L4_ is significantly higher. (**D**) Comparison between representational similarity of iEEG vs. fMRI and iEEG vs. DCNNs. Left, RS_fMRI-V1_ (blue) vs.RS_Raw-Resnet50-L1_ (black). Right, RS_fMRI-VOT_ (red) vs. RS_SHINE-Alexnet-L4_ (black). fMRI-V1 and Raw-Resnet50-L1 are the best low-order references for iEEG in respective modalities, whereas fMRI-VOT and SHINED-Alexnet-L5 are the best high-order references. Colored/black asterisks indicate when RS_fMRI_/RS_DCNN_ is significantly higher. The best-matching DCNN layer didn’t correspond to late LOC and VOT responses to the same extent as same-area fMRI responses.

**Fig. S4.**
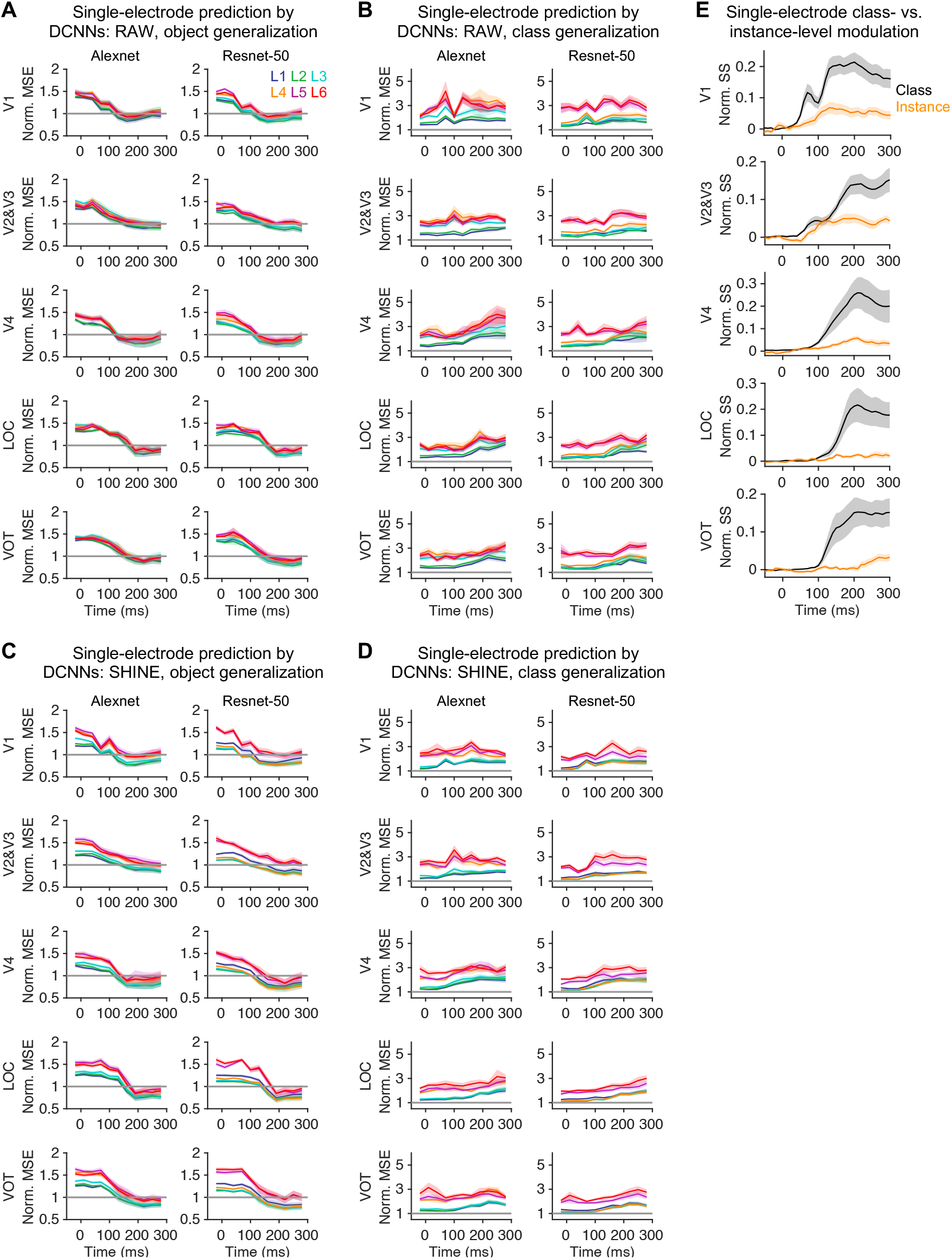
Predicting single-electrode responses with DCNN output. (**A**) Performance (crossvalidated normalized mean square error) of predicting single-electrode, instance-level responses with linearly mixed output of layers L1 to L6 of Alexnet (left) and Restnet-50 (right) using raw stimuli as input. We did his analysis for one of every three TOIs as it was computationally intensive. Lines and shadings indicate mean and standard error of mean over same-area electrodes. Training and test data were separate instances regardless of their class (object generalization). (**B**) As (A), the predictive performance of class generalization, in which training and test data were instances from separate classes. (**C** and **D**) As (A) and (B), respectively, predictive performance when SHINED stimuli served as input to DCNNs. DCNN-based models passed the object generalization test, which predicted singleelectrode responses better than baseline (norm. MSE < 1), i.e., a flat line through the mean, taking either raw or SHINED stimuli as input of DCNNs. However, the models failed the class generalization test (norm. MSE > 1), a test of model validity proposed in previous work^24^ that is stronger than object generalization. (**E**) Class- and instance-level modulation of single-electrode, single-trial responses, quantified as the bias-corrected sum of squares (SS) at the class and instance levels divided by SS of trial-by-trial errors, respectively (see Methods). Classes rather than instances (of the same classes) explained single-electrode modulation much better, especially in V4, LOC, and VOT, which suggests the inherent dimensionality of 120-dim instance-level responses might be no larger than 6 (# classes). Considering the predictive models for multiple electrodes in matrix form, i.e., Y (responses; # electrodes × # instances) = beta (mixing weights; # electrodes × # units plus a constant) × X (DCNN output; # units plus a constant × # instances), the rank of both response and weight matrices (X and beta) should also be six or fewer. Finding the weights then became trivial because the DCNN output (X) should be close to full rank (120) due to high # units (1k to 800k) for all layers and both models, which implies the predictive performance may not be a proper metric of brain-DCNN correspondence. Indeed, the relative performance (object generalization) of L1 through L6 was largely the same regardless of which area’s response to predict and at which time for the same model/input. The weights (beta) learned from instances of a subset of classes (class generalization) should have a lower rank than that learned from instances of all classes (object generalization), which may explain the poor predictive performance of the former that was indicative of severe overfitting during training. In a recent study^31^ that reported good DCNN-based predictive performance for fMRI responses, more repetitions per image and averaging over voxels and subjects may have increased the fidelity of instance-level responses. Such a strategy could, in the future, improve the feasibility/validity of assessing brain-DCNN correspondence from the response prediction approach.

**Fig. S5.**
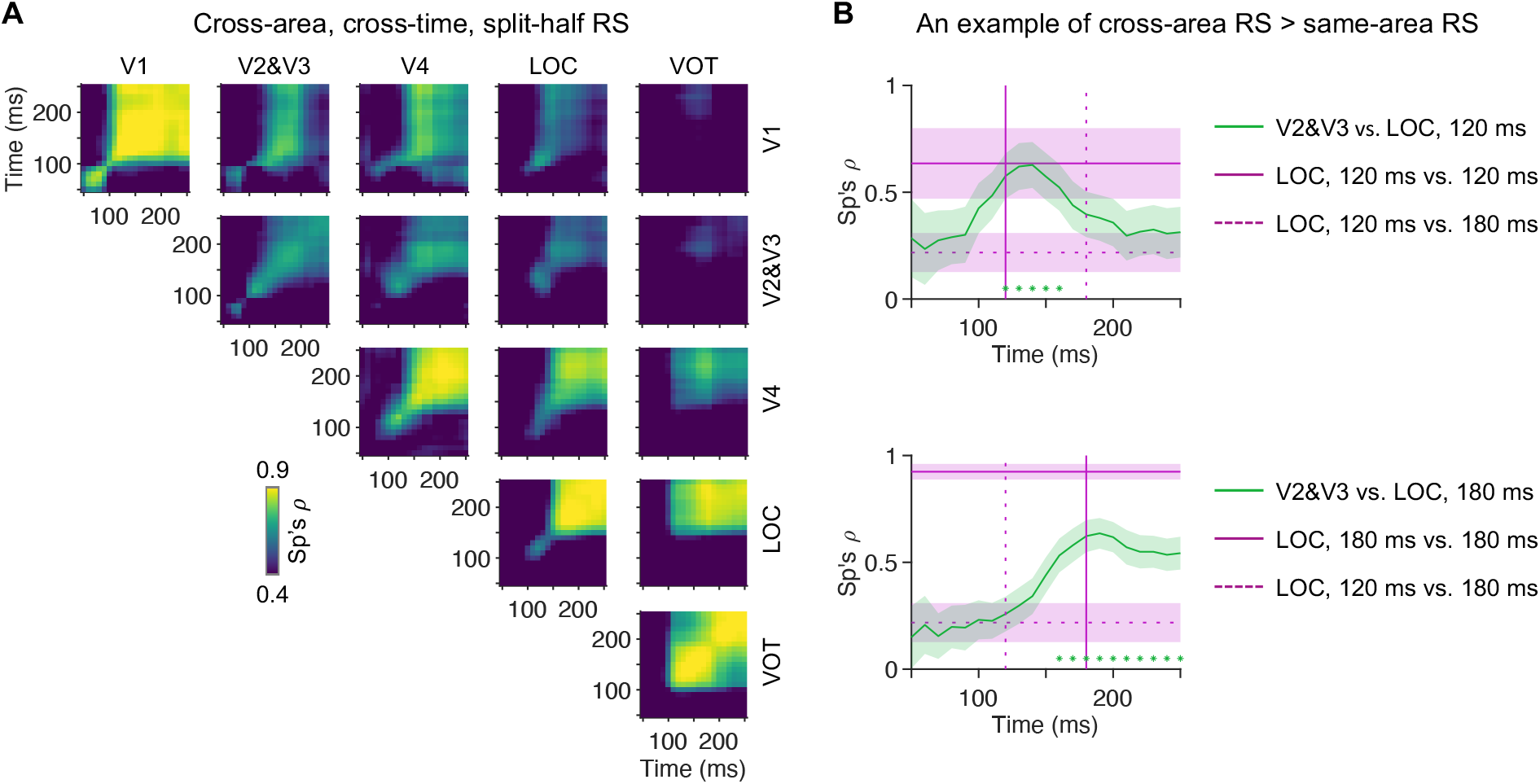
Relating neural representations across areas and time. (**A**) Cross-area, cross-time RS of inter-class RDMs built using separate trials (bootstrap resampling and split-half; see Methods). X- and Y-axes indicate the time of RDMs to be compared, which could be of the same or different areas. The mean value over bootstrap distribution is shown. (**B**) An example of higher cross-area, close-time RS than same-area, distant-time RS. Top, RS of V2&V3 at 50 to 250 ms vs. LOC at 120 ms (green), compared to RS of LOC at 120 ms vs. itself (purple solid) and LOC at 180 ms (purple dotted). The latter two are drawn as flat lines as they only correspond to one time point each. Lines and shadings indicate the mean and standard error of bootstrap distribution. Green asterisks indicate when RS of V2&V3 vs. LOC at 120 ms is significantly higher than RS of LOC at 120 ms vs. LOC at 180 ms (P < 0.05). Vertical solid/dotted line indicates 120/180 ms, i.e., corresponding to the respective flat line. Bottom, as top, RS of V2&V3 vs. LOC at 180 ms (green), compared to RS of LOC at 180 ms vs. itself (purple solid) and LOC at 120 ms (purple dotted).

**Fig. S6.**
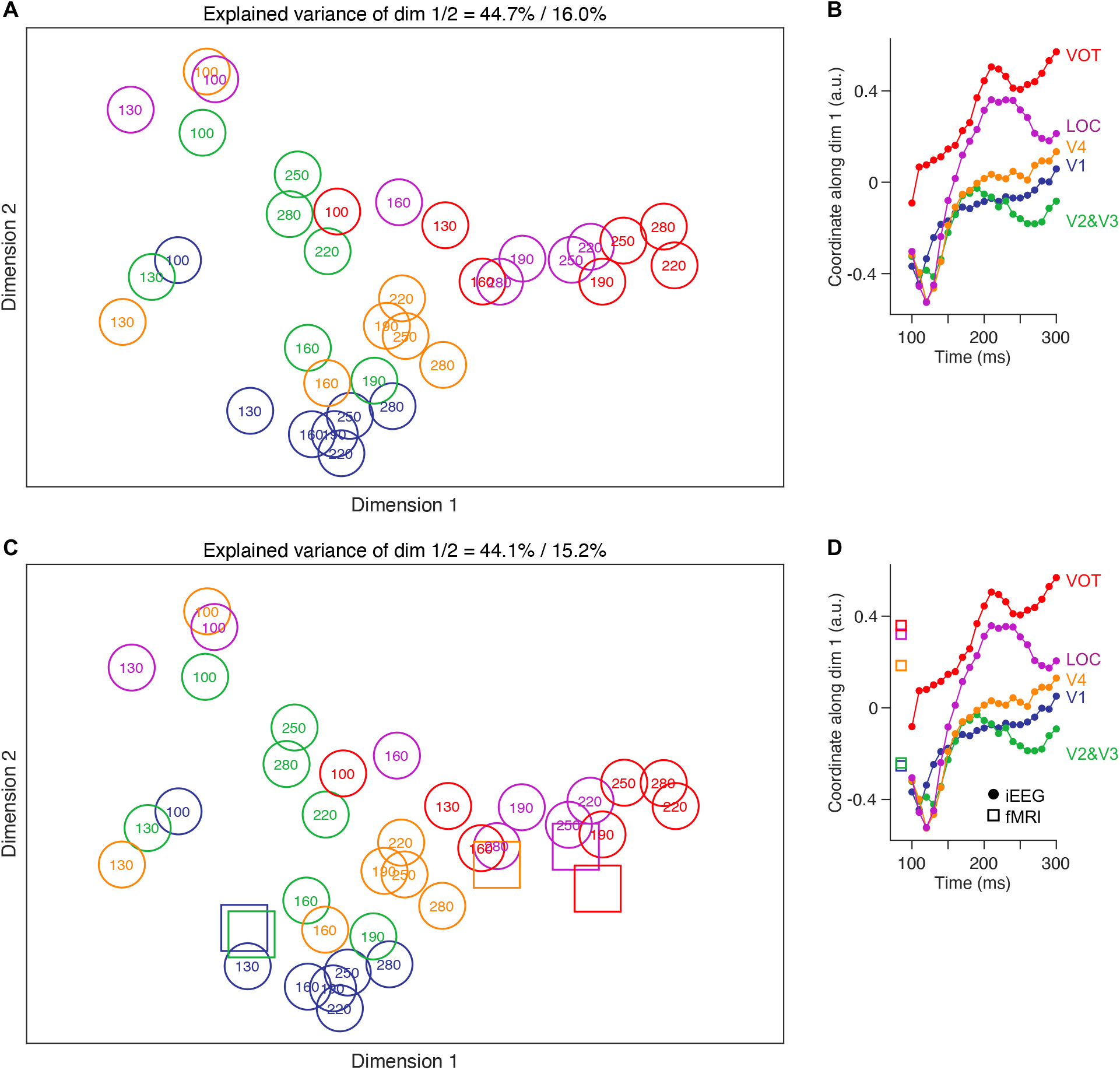
Multidimensional scaling of neural representations across areas and time. (**A**) MDS visualization of iEEG RDMs across areas and time. We applied classical MDS (*cmdscale* in MATLAB) to split-half RS between inter-class RDMs in five areas at 100 to 300 ms because this algorithm reports explained variation. Coordinate along the first two MDS dimensions of RDMs at one of every three TOIs is shown to avoid clutter. The color indicates the area, and the text indicates time (ms). (**B**) Coordinate along the first MDS dimension of RDMs at all TOIs from 100 to 300 ms is shown by area. The first dimension accounted for a much larger portion (44.7%) of variation than the others. The coordinate tended to be more positive for RDMs in higher areas and at later time points. Nonetheless, distance in MDS space would better approximate the full matrix of RS (see fig. S5A) by adding more dimensions. The results suggest representational changes across OTC and lower areas were largely aligned, although they were not strictly one-dimensional. (**C**) and (**D**), as (**A**) and (**B**), results of MDS applied to both iEEG and fMRI RDMs. The pattern didn’t change much as there were fewer fMRI RDMs (squares), which typically had their dim. one coordinate within the range of same-area iEEG RDMs, except V4. The latency of best-matching iEEG RDM for respective same-area fMRI RDM suggests recurrent processing contributes to fMRI responses in LOC and VOT.

